# Transient RNA structures cause aberrant influenza virus replication and innate immune activation

**DOI:** 10.1101/2022.01.25.476955

**Authors:** Hollie French, Emmanuelle Pitré, Michael S. Oade, Elizaveta Elshina, Karishma Bisht, Alannah King, David L.V. Bauer, Aartjan J.W. te Velthuis

## Abstract

During infection, the influenza A virus RNA polymerase produces both full-length and aberrant RNA molecules, such as defective viral genomes (DVG) and mini viral RNAs (mvRNA). Subsequent innate immune activation involves the binding of host pathogen receptor retinoic acid-inducible gene I (RIG-I) to viral RNAs. However, not all influenza A virus RNAs are strong RIG-I agonists. Here we show that potent innate immune activation by mvRNAs is determined by transient RNA structures, called template loops (t-loop) that stall the viral RNA polymerase. The effect of t-loops depends on the formation of an RNA duplex near the template entry and exit channels of the RNA polymerase, and their effect is enhanced by mutation of the template exit path from the RNA polymerase active site. Overall, these findings provide a mechanism that links aberrant viral replication to the activation of the innate immune response.

## Introduction

Influenza A viruses (IAV) are important human pathogens that generally cause a mild to moderately severe respiratory disease. A range of viral, host, and bacterial factors can influence the outcome of infections with IAV(*1, 2*). One important factor is the activation of host protein retinoic acid-inducible gene I (RIG-I) by double-stranded 5’ di- or triphosphorylated RNA(*3*). Activated RIG-I translocates to mitochondria and triggers oligomerization of mitochondrial antiviral signaling protein (MAVS), leading to the expression of innate immune genes, including interferon-β (IFN-β) and IFN-λ(*4-6*). Innate immune gene expression typically leads to a protective antiviral state, but results in an overproduction of cytokines and chemokines when dysregulated. This phenomenon underlies the lethal pathology of infections with 1918 H1N1 pandemic or highly pathogenic avian IAV(*7, 8*), and various viral and host factors have been implicated in causing immunopathology, including the products of aberrant viral replication(*9*).

During an IAV infection, the virus introduces eight ribonucleoproteins (RNP) into the host cell nucleus. These RNPs consist of oligomeric viral nucleoprotein (NP), a copy of the viral RNA polymerase, and one of the eight segments of single stranded negative sense viral RNA (vRNA) that make up the viral genome(*10*) (Fig. 1A). The vRNA segments range from 890 to 2341 nt in length, but all contain conserved 5’ triphosphorylated, partially complementary 5’ and 3’ termini(*10*). These termini serve as promoter for the RNA polymerase, but also as agonist of RIG-I(*3, 11*). In the context of an RNP, the termini are bound by the RNA polymerase subunits PB1, PB2, and PA(*12, 13*), likely to prevent RIG-I binding to the vRNA segments(*5*). During viral replication, a second RNA polymerase is recruited to the RNP to encapsidate nascent RNA(*14, 15*), suggesting that access of RIG-I to viral RNA is also impaired during active replication.

**Figure 1.**
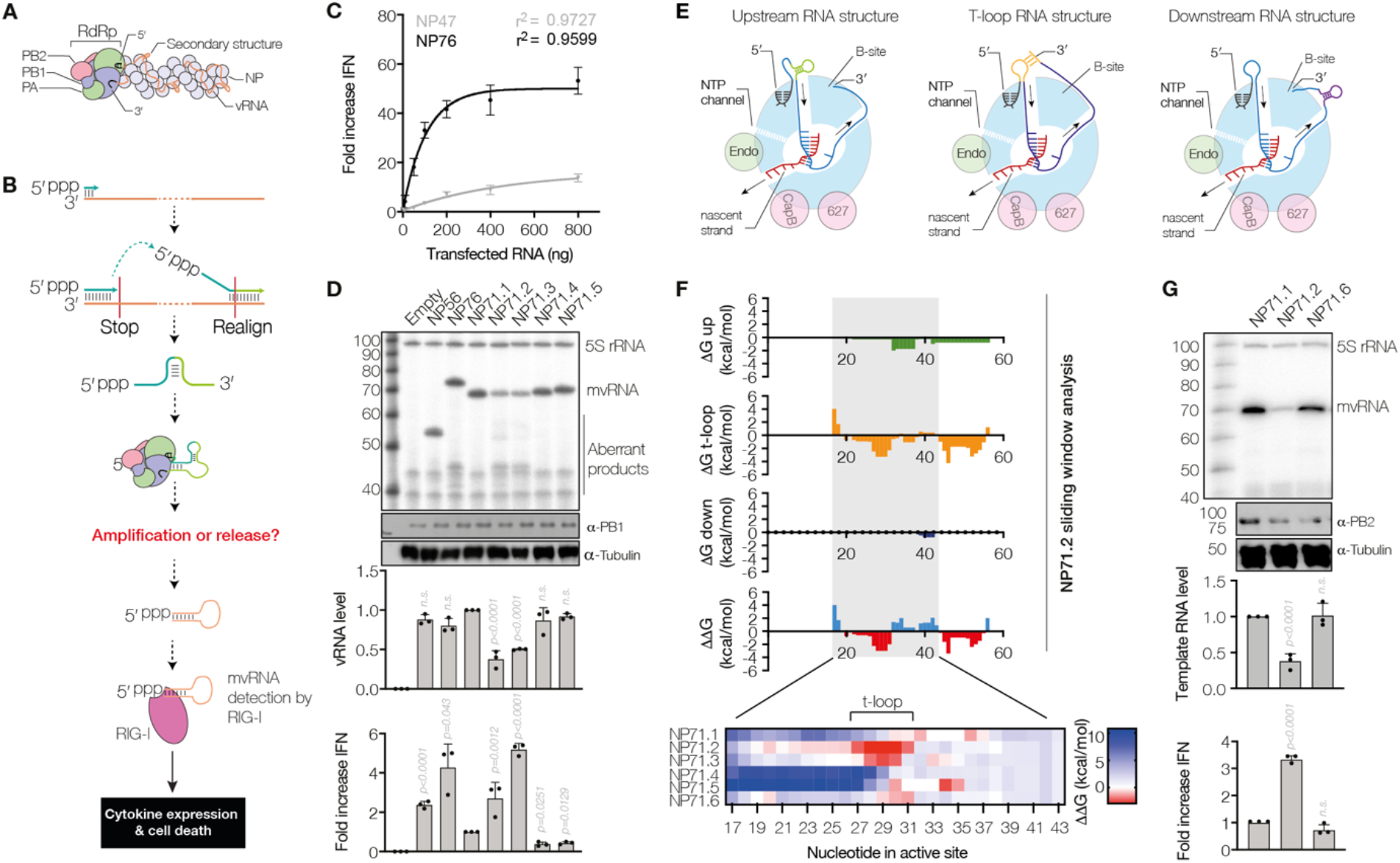
Sequence-dependent reduction of mvRNA replication and induction of IFN promoter activation. A) Schematic of influenza A virus RNP. B) Schematic of mvRNA formation via intramolecular template switching. This process has the potential to create novel RNA structures. C) IFN-β promoter activation following transfection of in vitro transcribed segment 5 mvRNAs of 47 or 76 nt in length. D) Replication of model mvRNAs in HEK 293T cells by the IAV A/WSN/33 (H1N1) RNA polymerase. RNA levels were analyzed by primer extension and the ability of mvRNA replication to induce IFN-β promoter activity was analyzed using a luciferase reporter assay. E) Schematic of RNA structure formation upstream, around (t-loop), and downstream of the RNA polymerase. F) ΔG values for RNA structures forming upstream (green), around (t-loop; orange) or downstream (purple) of the RNA polymerase were computed using a sliding window approach. The difference in ΔG (ΔΔG) between the formation of a t-loop and structures forming upstream or downstream the RNA polymerase was computed and shown in the bottom graph. Heatmap shows zoom in on ΔΔG values computed for middle of the mvRNA templates used. G) Replication of the NP71.1, NP71.2 or the destabilized NP71.6 mvRNA templates in HEK 293T cells by the WSN RNA polymerase. RNA levels were analyzed by primer extension and the ability of mvRNA replication to induce IFN promoter activity was analyzed using a luciferase reporter assay.

In addition to full-length vRNA and cRNA molecules, the viral RNA polymerase can produce RNAs that are shorter than the vRNA or cRNA template from which they derive. Such defective viral genomes (DVGs)(*16*) and mini viral RNAs (mvRNA)(*9*) are >180 nt and 56-125 nt long aberrant viral RNA molecules, respectively, that contain internal deletions between the conserved 5’ and 3’ termini(*11*). Among the different viral RNA species, RIG-I preferentially binds to and is most potently activated by mvRNAs(*9*). Moreover, the RNA polymerases of highly pathogenic avian H5N1 and pandemic 1918 H1N1 IAV produce higher mvRNA levels than the RNA polymerase of lab adapted H1N1 IAV(*9*), suggesting that there is a correlation between mvRNA production and innate immune activation in infections with highly pathogenic IAV. It is presently not known why mvRNAs are such potent agonists.

mvRNAs are generated, in part, via a copy-choice mechanism that results in the loss of an internal genome segment sequence(*9, 17*) (Fig. 1B). As a result, RNA sequences or structures that do not normally reside side-by-side in the full-length genome segments are brought closer to each other in the nascent RNA, potentially resulting in the formation novel RNA structures (Fig. 1B). Inherently, the RNA polymerase is not impaired by RNA structures in the RNP and it can replicate and transcribe an mvRNA containing a copy of the aptamer Spinach(*18*), a highly-structured RNA capable of stabilizing the fluorophore DFHBI (Fig. S1). However, it is not known if RNA secondary structures play a role in the activation of the innate immune response. Once generated, mvRNAs can be replicated by the viral RNA polymerase in the absence of NP(*19*). Because the viral RNA polymerase and RIG-I compete for binding to the same 5’ and 3’ termini, RIG-I activation can only be induced by mvRNAs that have dissociated from the RNA polymerase or that were exceptionally abundant and not bound by the RNA polymerase complex. Here we demonstrate that mvRNAs capable of inducing innate immune responses are defined by transient RNA structures that we call t-loops. These t-loops affect viral RNA polymerase processivity, and induce aberrant RNA synthesis and IFN-β promoter activation.

## Results

### Induction of IFN-β promoter activation by mvRNAs is sequence dependent

It is unclear what determines whether an influenza virus mvRNA is a strong inducer of the innate immune response. To systematically investigate if the sequence or secondary structure of an mvRNA template can affect IAV RNA polymerase activity, we engineered five segment 5-derived mvRNA templates. Each engineered mvRNA had a length of 71 nt (NP71.1-NP71.5), but a different internal sequence (Table S1). Our positive control mvRNAs were previously described 56- and 76-nt long mvRNAs derived from segment 5 (NP56 and NP76, respectively), and our negative control mvRNA was a 47-nt long mvRNA derived from segment 5 (NP47) that is unable to bind RIG-I and induce a strong IFN signal(*9*). Indeed, transfection of increasing amounts of in vitro transcribed NP76 into HEK 293T cells resulted in a strong exponential increase in IFN-β promoter activity (λ = 74 ng), while the NP47 induced a much-reduced activity (λ = 190 ng) (Fig. 1C), showing how these RNAs differentially induce IFN-β promoter activity when are transfected into the cytoplasm.

We subsequently investigated the replication of the 71-nt long mvRNA templates by the IAV RNA polymerase. To this end, we transfected plasmids expressing the viral RNA polymerase subunits PB1, PB2 and PA, a plasmid expressing NP, and a plasmid expressing a template mvRNA into HEK 293T cells. Primer extension analysis showed efficient amplification of NP76 as well as the production of several smaller aberrant RNA products <76 nt in length (Fig. 1D). Fractionation of cells in which NP76 was replicated showed that the NP76 mvRNA template was present in the nuclear, cytoplasmic, and mitochondrial fractions, whereas the aberrant RNAs produced from NP76 were present in the nucleus only (Fig. S2). Since IAV RNA is predominantly detected in the cytoplasm of the host cell, these results suggest that only the mvRNA template, and not aberrant products shorter than the template mvRNA, play a role in innate immune activation. Subsequent analysis of the engineered mvRNA templates showed that three of these templates were efficiently replicated (NP71.1, NP71.4 and NP71.5), while the other two mvRNAs (NP71.2 and NP71.3) were not (Fig. 1D). Interestingly, among the engineered mvRNAs, templates that were poorly replicated showed higher IFN-β promoter activity and aberrant RNA synthesis (i.e., the production of RNA products shorter than the template) than the three mvRNA templates that were efficiently replicated (Fig. 1D).

To exclude that a differential recognition of the engineered mvRNAs by the host cell was responsible for the observed increased IFN-β promoter activity on the NP71.2 and NP71.3 mvRNAs, we isolated total RNA from HEK 293T transfections and re-transfected equal amounts of these RNA extracts together with IFN-β and *Renilla* reporter plasmids into HEK 293T cells. Re-transfection of NP71.1-NP71.5 showed an inverse pattern of IFN-β promoter activation in comparison to Fig. 1D, whereby abundant mvRNAs induced more IFN-β promoter activity than the least abundant mvRNAs (Fig. S3), suggesting that there is no inherent difference between the mvRNA in their ability to activate IFN-β promoter activity. Instead, these results indicated that active viral replication determines whether an mvRNA will activate innate immune signaling in the context of an RNP.

To verify that the different replication efficiencies had not been the result of the effect of NP71.2 and NP71.3 on the innate immune response, we also expressed these mvRNAs and the WSN RNA polymerase in *MAVS*^−/−^ IFN::luc HEK 293 cells(*20*). These *MAVS*^−/−^ cells do not express endogenous MAVS (Fig. S4A), blocking any RIG-I mediated innate immune signaling, but overexpression of a MAVS-FLAG plasmid still triggers IFN-β promoter activity, indicating that the IFN-β reporter is still functional (Fig. S4B). Expression of NP71.1-NP71.5 in the *MAVS*^−/−^ cells did not induce IFN-β promoter activity (Fig. S4C). Subsequent primer extension analysis showed that the differences in replication between NP71.1-NP71.5 had been maintained in the *MAVS*^−/−^ cells (Fig. S4C), demonstrating that the differential replication efficiency is not dependent on the innate immune response.

To investigate whether the effect of the NP71.3 and NP71.4 mvRNAs was specific to the WSN polymerase, we next expressed these mvRNAs alongside the pandemic H1N1 A/Brevig Mission/1/18 (abbreviated as BM18) or the highly pathogenic avian H5N1 A/duck/Fujian/01/02 (abbreviated as FJ02) RNA polymerases. We found that the BM18 and FJ02 RNA polymerases were impaired on the NP71.2 and NP71.3 mvRNA templates and triggered a stronger IFN-β promoter activity on the NP71.3 template relative to the NP71.1 template (Fig. S5). We also observed that the BM18 RNA polymerase produced short aberrant RNA products, while the FJ02 RNA polymerase did not, despite inducing IFN-β promoter activity (Fig. S5). Together, these results suggest that the mvRNA template is the innate immune agonist, rather than the aberrant RNA products derived from the template, and that innate immune activation is dependent on a sequence-specific interaction between the active IAV RNA polymerase and the mvRNA template.

### T-loops affect viral polymerase activity and IFN-β promoter activation

The viral RNA template enters and leaves the active site of the IAV RNA polymerase as a single-strand through the entry and exit channels, respectively (Fig. 1E)(*21-23*). However, the IAV genome contains various RNA structures that need to be unwound(*24*). Moreover, unwinding of these structures may lead to the formation of transient RNA structures upstream or downstream of the RNA polymerase that may modulate RNA polymerase activity (Fig. 1E), while base pairing between a part of the template that is entering the RNA polymerase and a part of the template that has just been duplicated may trap the RNA polymerase in a template loop (t-loop) (Fig. 1E). To systematically analyze what (transient) RNA structures are present during replication, we used a sliding-window algorithm to calculate the minimum free energy (ΔG) for every putative t-loop as well as every putative secondary RNA structure upstream and downstream of the RNA polymerase (Fig. S6A-B). For each position analyzed, we excluded 20 nt from the folding analysis for the footprint of the IAV RNA polymerase(*21*) and 12 nt from the 5′ terminus, which is stably bound by the RNA polymerase prior to replication termination. As shown in Fig. 1F and Fig. S6C, our analysis revealed that NP71.2 and NP71.3 are unique among the engineered mvRNA templates in forming t-loop structures around nucleotide 29 of the positive sense replicative intermediate (cRNA), but not the negative sense (Fig. S6D), suggesting that t-loops in the positive sense mvRNA template modulate RNA polymerase activity. The likelihood that the t-loops form in the context of other secondary structures was calculated as the difference (ΔΔG) between the computed ΔG values for the individual structures (Fig. 1F).

To confirm that t-loops affect RNA polymerase processivity and IFN-β promoter activity, we replaced two A-U base pairs of the NP71.2 t-loop duplex with two G-U base pairs, creating NP71.6 (Fig. 1G, S7A). Using our sliding window analysis, we confirmed that this mutation would make t-loop formation near nucleotide 29 less favorable (Fig. 1F). Following expression of NP71.1, NP71.2 and NP71.6, we found that replication of the NP71.6 mvRNA was increased relative to the NP71.2 mvRNA, our control mvRNAs, and the NP71.1 mvRNA (Fig. 1G, S7B). In addition, destabilization of the t-loop reduced the induction of the IFN-β promoter activity (Fig. 1G). By contrast, when we mutated the stem of the t-loop of the NP71.2 mvRNA template such that the t-loop around nucleotide 29 was maintained (Fig. S7C; NP71.7-8, replication remained reduced and IFN-β reporter activity increased relative to the NP71.1 mvRNA (Fig. S7C-D). We also observed that replication of all mvRNAs with a t-loop led to the production of short aberrant RNA products. However, increases in aberrant RNA levels were not correlated with increases in IFN-β reporter activity, in line with the results in Fig. 1C and S1, and indicated that the mvRNA template is the agonist of IFN-β reporter activity. Together these results show that t-loops negatively affect IAV RNA synthesis and stimulate innate immune signaling during IAV replication.

### T-loops reduce RNA polymerase processivity in vitro

The mvRNAs NP71.2 and NP71.3 contain a t-loop in the first half of the positive sense mvRNA template. To confirm that t-loops also affect RNA polymerase activity in the negative sense, we engineered three additional 71-nt long mvRNA templates with t-loops in different locations of the template (NP71.10-12) (Table S1; Fig. 2A). Expression of these mvRNA templates together with the subunits of the viral RNA polymerase in HEK 293T cells led to strongly reduced NP71.10 and NP71.11 mvRNA levels and partially reduced NP71.12 mvRNA levels (Fig. 2B). In line with our other results (Fig. 1), IFN-β promoter activity was increased for the NP71.11 and NP71.12 templates relative to the NP71.1 mvRNA, while the NP71.10 mvRNA did not induce IFN-β promoter activity, likely because it was too poorly or not fully replicated (Fig. 2B).

**Figure 2.**
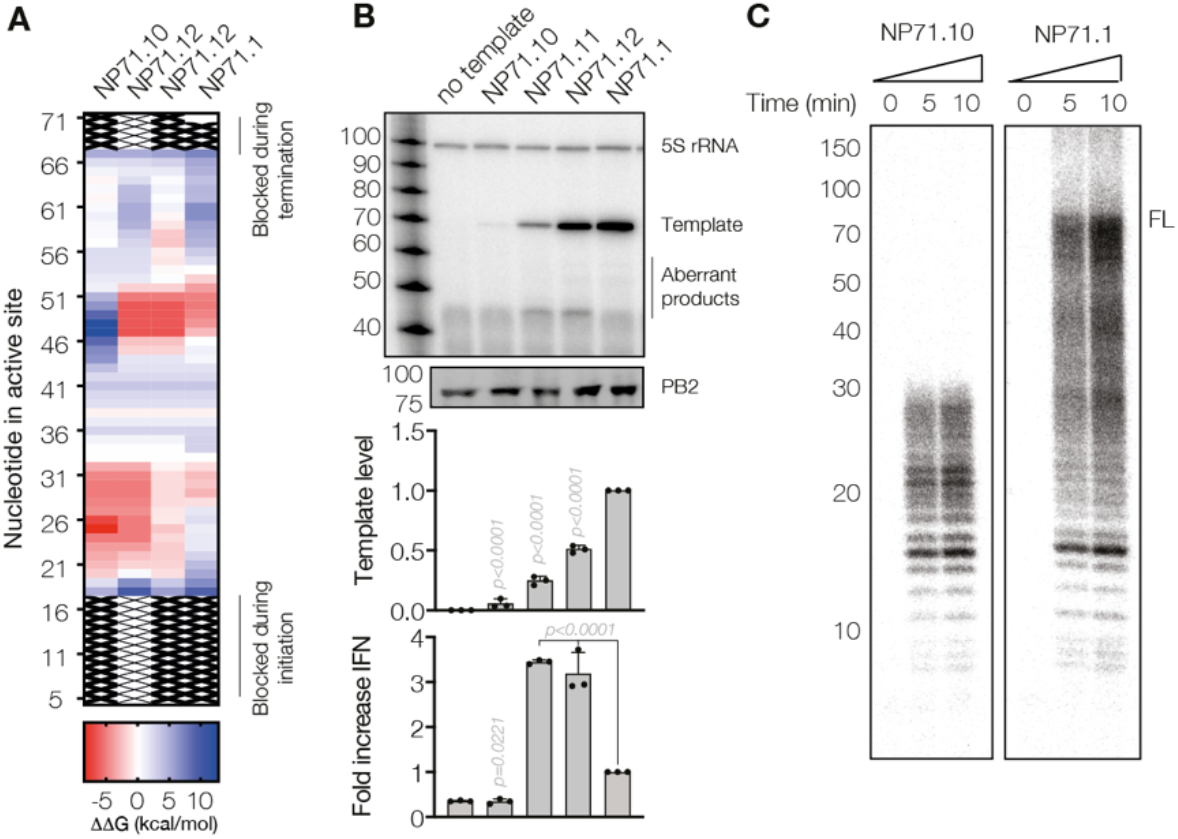
mvRNA t-loops stall RNA polymerase activity and induce IFN-β promoter activation. A) Heat map showing ΔΔG as estimate for the likelihood of t-loop formation. B) Replication of mvRNA templates. Top panel shows primer extension analysis. Graphs show quantification of template RNA level and IFN-β promoter activation. C) Analysis of IAV RNA polymerase activity in vitro.

To investigate if t-loops affect RNA polymerase processivity *in vitro*, we purified the WSN RNA polymerase from HEK 293T cells using tandem-affinity purification (TAP)(*25*) and incubated the enzyme with the NP71.1 and NP71.10 mvRNA templates in the presence of NTPs. Following denaturing PAGE and autoradiography, we observed a main product of approximately 71 nt in reactions containing the NP71.1 control mvRNA (Fig. 2C, S8A). By contrast, incubations with the NP71.10 mvRNA templates resulted in products up to approximately 27 nt in length, in agreement with the location of the t-loop in the first half of the mvRNA template (Fig. 2A). Moreover, this result offered a possible explanation for the lack of IFN-β promoter activity induction by the NP71.10 template (Fig. 2B).

To investigate if the effect of the t-loops could be modulated by temperature, we expressed the mvRNA templates together with the viral RNA polymerase in HEK 293T cells grown at 37 °C or 39 °C. After 24 hours, we observed a reduction in the replication of the mvRNA templates at 39 °C relative to 37 °C, and concomitant reduction in IFN-β promoter activity (Fig. S8B). This result is in line with previous observations that at higher temperatures IAV replication is impaired(*26*). However, the reduction in template mvRNA levels was more reduced for the NP71.11 relative to the NP71.1 mvRNA (Fig. S8C), suggesting that destabilization of the t-loop in the NP71.11 template had increased viral replication relative to the NP71.1 mvRNA template. Together these observations indicate that t-loops affect influenza A virus polymerase processivity on mvRNAs and that reduced RNA polymerase processivity underlies IFN-β promoter activation.

### PB1 K669A increases t-loop sensitivity and IFN-β promoter activation

In the IAV RNA polymerase elongation complex, the 3’ terminus of the template is guided out of the template exit channel via an exit groove on the outside of the thumb subdomain. This groove consists of PB1 and PB2 residues, and leads to promoter binding site B(*13, 22*) (Fig. 1E, 3A). Since this exit grove and the template entry channel reside next to each other at the top of the RNA polymerase (Fig. 1E, 3A), perturbation of the path of the 3’ terminus out of the exit channel may lead to t-loop formation, reduced RNA polymerase activity, and increased IFN-β promoter activation (Fig. 1E, 3A). To test if dysregulation of the exit groove leads to more IFN-β promoter activation, we mutated PB1 lysine 699, which resides at the start of the exit groove (Fig. 3B), to alanine (K669A). Mutation of this residue had no effect on RNA polymerase activity in the presence of a full-length segment 6 template(*27*) (Fig. 3C) or the NP71.1 and NP71.6 mvRNA templates that do not contain a stable t-loop (Fig. 3D-E). However, when we expressed the K669A mutant together with the NP71.2, NP71.11 or NP71.12 mvRNAs, which do contain a t-loop in either the positive or negative sense, the K699A mutant displayed reduced RNA polymerase activity (Fig. 3D-E). Interestingly, on all templates, the K699A mutant induced significantly higher IFN-β promoter activity relative to the wild-type RNA polymerase, and considerably more on the t-loop containing templates (Fig. 3B-E). These results thus imply that mutation of the exit groove increases the base-level potential of the RNA polymerase to trigger FN-β promoter activity, and that it makes the RNA polymerase more sensitive to disruption by a t-loop than the wildtype RNA polymerase, establishing a link between polymerase structure, RNA polymerase dysregulation, and innate immune activation.

**Figure 3.**
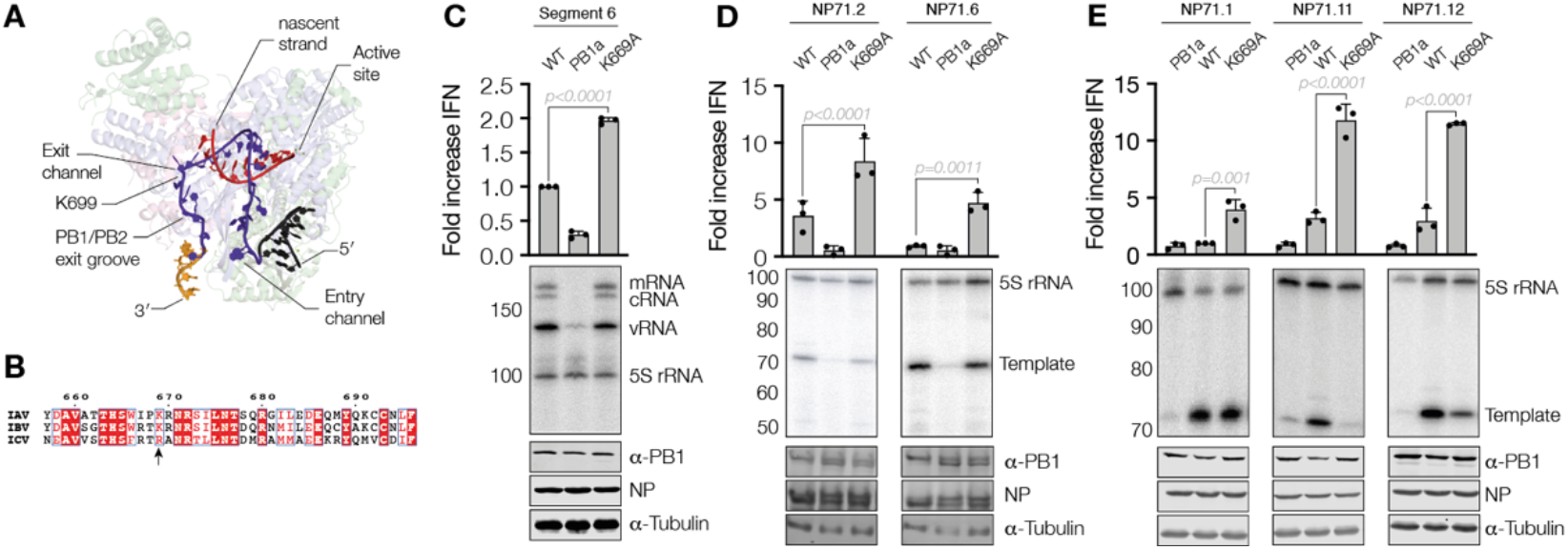
Mutation near template exit channel increases RNA polymerase sensitivity to t-loops. A) Structure of the pre-termination complex of the bat influenza A virus RNA polymerase (PDB 6SZU). The 5’ and 3’ ends of the template are shown in black and gold, respectively. The body of the template is shown in dark blue and the nascent RNA in red. Location of PB1 K669 is indicated. B) Amino acid alignment of PB1 C-terminus. PB1 K669 is indicated with an arrow. C) IFN-β promoter activity and segment 6 viral RNA levels in the presence of wildtype and K669A RNA polymerases. For the segment 6 vRNA template, the mRNA, cRNA and vRNA species are indicated. 5S rRNA and western blot for PB1 subunit expression are shown as loading control. D-E) IFN-β promoter activity and mvRNA template levels for five NP71 mvRNA templates in the presence of wildtype and K669A RNA polymerases. RNA levels were analyzed by primer extension. 5S rRNA and western blot for PB1 subunit, NP, and tubulin expression are shown as loading control.

### IFN-β promoter activation by mvRNAs during infection is t-loop dependent

mvRNAs are produced during IAV infection *in vitro* and *in vivo*(*9*). To study how their sequence and abundance varies, we analyzed RNA extracted from ferret lungs 1 day post infection with BM18 for 1 day and A549 cells infected with WSN for 8 hours. Although no quantitative comparisons can be made due to the different infection conditions, we do find a strikingly similar variation in mvRNA sequence and abundance (Fig. 4A-B).

**Figure 4.**
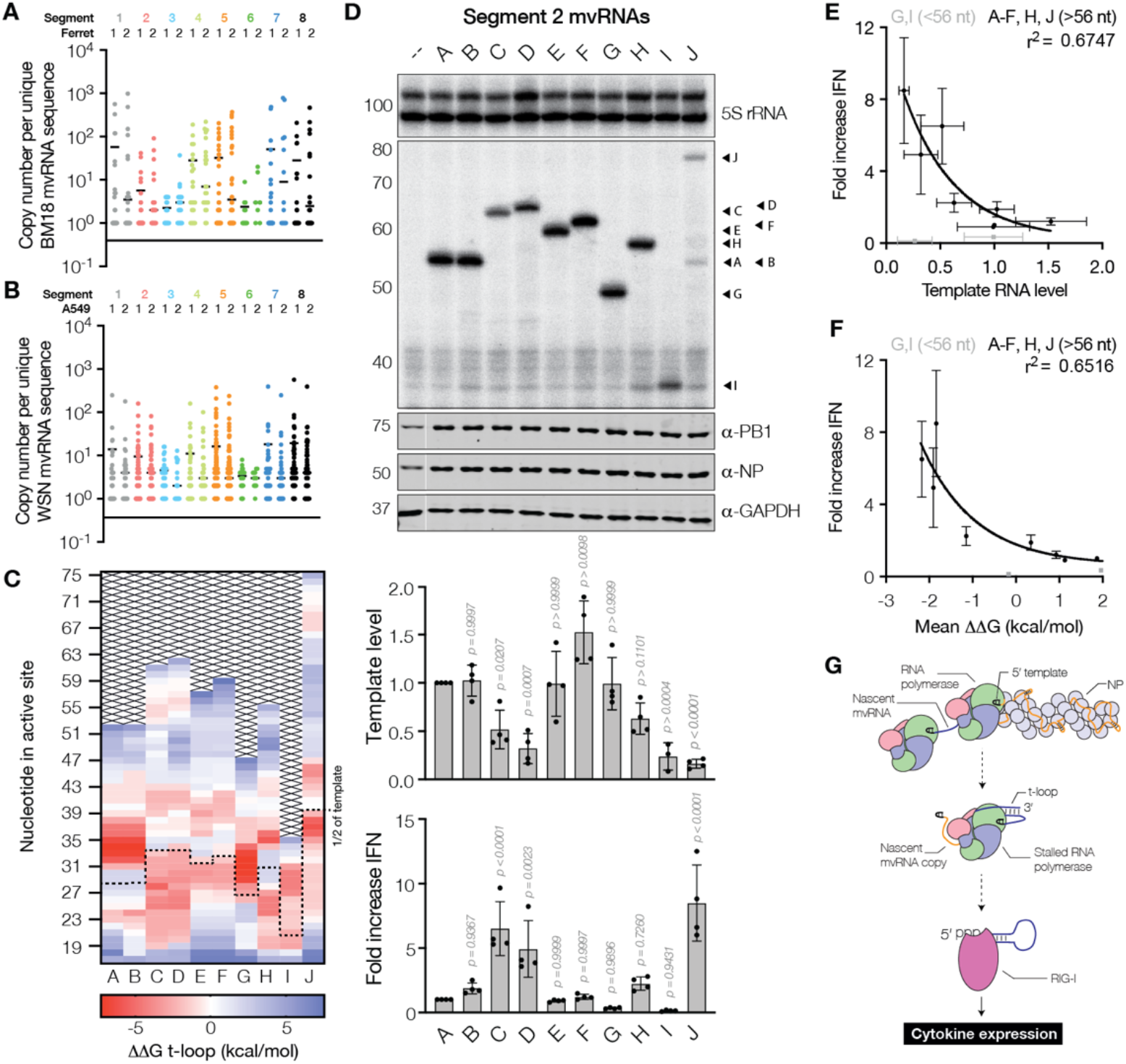
Reduced mvRNA replication in viral infections is correlated with IFN-β promoter activation. A) Number of mvRNA copies per unique mvRNA sequence detected in ferret lungs infected with BM18 or B) A549 cells infected with WSN. In each graph two biological repeats are shown. C) ΔΔG heat map of negative sense template of WSN segment 2 derived mvRNAs. Half of the mvRNA template is indicated with a dotted line. D) Replication of segment 2-derived mvRNAs identified by NGS in HEK 293T cells by the WSN RNA polymerase. RNA levels were analyzed by primer extension. The ability of mvRNA replication to induce IFN-β promoter activity was analyzed using a luciferase reporter assay. E) IFN-β induction is negatively correlated with template replication level. PB1-mvRNA G and I were excluded from the fit to the exponential decay, because they are shorter than the IFN-β promoter induction cut-off of 56 nt. F) IFN-β induction is negatively correlated with ΔΔG of first half of template mvRNA. G) Schematic of aberrant RNA synthesis by the IAV RNA polymerase. T-loops present in some mvRNAs lead to reduced RNA polymerase processivity. This may induce template release and/or binding of the mvRNA template to RIG-I. Host factor ANP32A, which plays a key role during cRNA and vRNA synthesis, is not shown for clarity(*14, 35*).

To investigate the implications of these mvRNA differences on the activation of the IFN-β promoter, we cloned ten WSN segment 2 mvRNAs into pPolI plasmids (mvRNAs A-J; Fig. S9A and Table S2). Computation of the ΔΔG revealed potential t-loops in the first half of the mvRNA sequences for mvRNAs C, D, H and J, and potential t-loops in the second half of the mvRNA templates for mvRNAs E, F and G (Fig. 4C). Subsequent expression of the authentic WSN mvRNAs alongside the WSN RNA polymerase in HEK 293T cells showed significant differences in mvRNA amplification (Fig. 4D). These differences were correlated with the abundancy detected by next generation sequencing (NGS) for seven of the cloned mvRNAs (Fig. S9B). In addition, we observed that replication of mvRNAs C, D and J leads to the appearance of products shorter than the template mvRNA (Fig. 4D), and that the appearance of these products is correlated with a reduced replication of the template mvRNA, in line with our findings in Fig 1.

To investigate if the different segment 2 mvRNA levels influenced the innate immune response, we measured the IFN-β promoter activity. We found that IFN-β promoter activity varied greatly, with mvRNAs C, D and J inducing the strongest response (Fig. 4D). Templates I and G, the two shortest mvRNAs at 52 and 40 nt long, induced the lowest IFN-β promoter activity, in line with our previous observations that mvRNAs <56 nt do not stimulate RIG-I and Fig. 1C. With mvRNAs I and G excluded due to their short size, these observations indicate that the IFN-β promoter activity is negatively correlated with mvRNA template level for mvRNAs >56 nt (Fig. 4E). Moreover, in line with our hypothesis presented in Fig. 1 that t-loops in the first half of the template affect RNA polymerase processivity, the IFN-β promoter activity was negatively correlated with the mean ΔΔG of the first half of the template (Fig. 4F). Weaker correlations were observed between the mvRNA length and the IFN-β induction, or the mvRNA length and mvRNA replication (Fig. S10A and B).

To exclude that a differential recognition of the mvRNAs was responsible for the observed anti-correlation, we isolated total RNA from HEK 293T transfections and retransfected equal amounts of these RNA extracts into HEK 293T cells. As shown in Fig. S11A, we observed no significant difference among the segment 2 mvRNAs longer than 56 nt. The mvRNAs G and I failed to induce a strong response due to their short length. To exclude that the different mvRNA levels had been the result of their different effects on the innate immune response, we also expressed the segment 2 mvRNAs in *MAVS*^−/−^ IFN::luc HEK 293 cells(*20*). Following expression of the segment 2 mvRNAs, we observed no IFN-β promoter activity (Fig. S11B). Primer extension analysis showed no significant reduction in mvRNA steady-state levels compared to wildtype cells (Fig. S11C), indicating that the replication of authentic mvRNAs is not impacted by innate immune activation. Together, these results confirm that the ability of a IAV mvRNA to induce IFN-β promoter activity is anti-correlated with the ability of the viral RNA polymerase to efficiently amplify it.

## Discussion

Two key factors important for the innate immune response in IAV infections are active viral replication and the binding of viral RNA molecules to RIG-I(*5*). RIG-I preferentially binds IAV mvRNAs over other viral RNA molecules(*9*), but, like full-length RNA molecules, mvRNAs are replicated by the viral RNA polymerase(*19*), which precludes access of RIG-I to these agonists. Here we show that impeded viral RNA polymerase processivity by t-loops determines the activation of innate immune signaling by mvRNAs. In mvRNAs, t-loops form when the 3’ terminus of the template can hybridize with an upstream part of the template (Fig. 1E). While the RNA polymerase may unwind a single t-loop, the formation of several successive t-loops in the first half of the mvRNA stalls the RNA polymerase. It is presently still unclear why a strong correlation is observed with t-loops in the first half of the template and not with downstream t-loops or how RIG-I gains access to the t-loop containing mvRNA once the polymerase has stalled (Fig. 4G). It is possible that, as shown previously for IAV polyadenylation(*22*), RNA polymerase stalling can result in an opening up of the RNA polymerase (defined as “skipping rope movement”). In turn, this may lead to release of the RNA template from the active site(*22*), potentially explaining why we observed mvRNA template accumulation in the cytoplasm and mitochondria, but no accumulation of more efficiently replicated aberrant RNA products (Fig. S2). Alternatively, nuclear RIG-I may directly interact with the stalled RNA polymerase complex. In contrast, the presence of a single t-loop with a long stable duplex or too many t-loops may prevent replication of the mvRNA and innate immune activation entirely (e.g., NP71.10).

It is possible that t-loops affect influenza RNA polymerase activity on full-length viral RNAs or DVGs. However, it is more likely that t-loops form only on partially formed RNPs. During viral RNA synthesis, NP dissociates and binds viral RNA in a manner that is coordinated by the viral RNA polymerase. When NP levels are reduced, aberrant RNPs or NP-less RNA products may form in which secondary RNA structures that are absent in the presence of NP contribute to t-loop formation RNA polymerase stalling(*24*). Indeed, this model can easily explain how reduced viral NP levels stimulate aberrant RNA synthesis and innate immune activation(*9, 28, 29*). We also observe that a mutation near the template exit channel increases the RNA polymerase sensitivity to t-loops (Fig. 3). Interestingly, we previously observed that avian adaptive mutations found near the template exit channel of highly pathogenic IAV RNA polymerases trigger mvRNA synthesis(*9*). It is thus tempting to speculate that the mechanism of mvRNA synthesis may involve RNA polymerase stalling by t-loops and we will address this possibility in future research.

During viral infections, mvRNA molecules of various lengths and abundancies are produced(*9*). We find that mvRNA abundance is not a perfect estimate for innate immune activation. Instead, we propose that mvRNAs that are poorly replicated contribute most to the activation of the innate immune system. By computing the ΔΔG based on primary sequence information, mvRNAs can be identified that trigger the innate response. Moreover, the link between transient RNA structure, RNA polymerase stalling, and innate immune activation that we reveal here may help explain conflicting observations in the literature what type of influenza virus RNA is a more potent RIG-I agonist. Instead, we suggest that the potent agonist may be unified by their ability to frustrate the processivity of the RNA polymerase on viral RNA molecules.

## Acknowledgements

The authors would like to thank Dr J. Rehwinkel for the HEK293 IFN:Luc and MAVS^−/−^ cells, Dr P. Palese for the A/Brevig Mission/1/18 (H1N1) polymerase subunit expression plasmids, and Dr Ian Goodfellow and Dr Aminu Jahun for the MAVS-FLAG expression plasmid. We thank Dr I. Olendraite, C. Rigby, Dr M. Richard, and Dr M. Funk for helpful discussions.

## Material and methods

### Viral protein and RNA expression plasmids

pcDNA3-based plasmids expressing influenza A/WSN/33 (H1N1) proteins PB1, PB2 PA, NP, PB2-TAP and the active site mutant PB1a (D445/D446A) have been described previously(*9, 30, 31*). Mutation K669A was introduced into the pcDNA3-PB1 plasmid by site-directed mutagenesis. mvRNA templates were expressed under the control of the cellular RNA polymerase I promoter from pPolI plasmids. PB1 mvRNA templates were generated by site-directed mutagenesis PCR deletion of pPolI-PB1. Short vRNA templates were created based on the pPolI-NP47 plasmid using the *Spe*I restriction site.

### Luciferase assay

Firefly luciferase reporter plasmid under the control of the *IFNB* promoter (pIFΔ(−116)lucter) and constitutively expressing *Renilla* luciferase plasmid (pcDNA3-*Renilla*) were described previously(*9*). The MAVS-FLAG expression vector and corresponding empty vector were cloned based on the pFS420, using the MAVS WT plasmid (pEF-HA-MAVS)(*32*).

#### Cells and transfections

Human embryonic kidney (HEK) 293T, Madin-Darby Canine Kidney (MDCK), and A549 cells were originally sourced from the American Type Culture Collection. All cells were routinely screened for mycoplasma. HEK293 wild-type and MAVS^−/−^ cells expressing luciferase under the control of the *IFNB* promoter were a kind gift from Dr J. Rehwinkel and were described previously(*20*). All cell cultures were grown in Dulbecco’s Modified Eagle Medium (DMEM) (Sigma) + 10% fetal bovine serum (FBS) (Sigma) and 1% L-Glutamine (Sigma). Transfections of HEK293T or HEK293 cell suspensions were performed using Lipofectamine 2000 (Invitrogen) and Optimem (Invitrogen) following the manufacturer’s instructions, and transfection of confluent, adherent HEK 293T cells were performed using PEI (Sigma) and Opti-Mem (Invitrogen).

#### Antibodies and western blotting

IAV proteins were detected using rabbit polyclonal antibodies anti-PB1 (GTX125923; GeneTex), anti-PB2 (GTX125926; GeneTex), and anti-NP (GTX125989; GeneTex) diluted 1:1000 in TBSTM (TBS/0.1% Tween-20 (Sigma)/5% milk). Cellular proteins were detected using the rabbit polyclonal antibodies anti-GAPDH (GTX100118; GeneTex) diluted 1:4000 in TBSTM, and anti-RNA Pol II (ab5131; Abcam) diluted 1:100 in TBSTM; the mouse monoclonal antibodies anti-MAVS E-3 (sc-166583; Santa Cruz) diluted 1:200 in TMSTM, and Mito tracker [113-1] (ab92824; Abcam) diluted 1:1000 TBSTM; and the rat monoclonal antibody anti-tubulin (MCA77G; Bio-Rad) diluted 1:1000 in TBSTM. Mouse monoclonal antibody anti-FLAG M2 (F3165; Sigma) diluted at 1:2000 was used to detect MAVS-FLAG. Secondary antibodies IRDye 800 Donkey anti-rabbit (926-32213; Li-cor), IRDye 800 Goat anti-mouse (926-32210; Li-cor), IRDye 680 Goat anti-mouse (926-68020; Li-cor), and IRDye 680 Goat anti-rat (926-68076; Li-cor), were used to detect western signals with a Li-cor Odyssey scanner.

#### RNP reconstitution assays and RNA sequence analysis

Infections and RNA analyses using primer extensions were performed as described previously(*9, 33*). mvRNA identification from next generation sequencing data was essentially performed as described previously(*9*), using data deposited in the Sequence Read Archive under accession number SUB3758924. T-loop analysis was performed using a custom Python script. Briefly, 20 nt of the template sequence were blocked off to represent the footprint of the viral RNA polymerase. This footprint was then moved in 1 nt increments along the template (Fig. S6A-B). T-loop formation was assessed by computing the ΔG of duplex formation between a stretch of 10 nt upstream of the footprint and 10 nt downstream of the footprint. The formation of upstream and downstream structures was computed for 24 nt windows upstream and downstream of the moving footprint. The ΔΔG was computed by subtracting The ViennaRNA package commands duplex-fold and cofold were used to compute the ΔG values(*34*).

#### Luciferase-based IFN expression assays

To measure IFN expression in RNP reconstituted HEK293T or HEK293 cells, luciferase assays were carried out 24 h post-transfection. RNP reconstitutions were carried out in a 24-well format by transfecting 0.25 µg of the plasmids pcDNA3-PB1, pcDNA3-PB2, pcDNA3-PA, pcDNA3-NP and a pPolI plasmid expressing a mvRNA template. HEK293T and HEK293 cells were additionally co-transfected with 100ng of the plasmids pIFΔ(−116)lucter and pcDNA3-*Renilla*. Cells were harvested in PBS and resuspended in an equal volume of Dual-Glo Reagent (Promega) followed by Dual-Glo Stop & Glo reagent (Promega). Firefly and *Renilla* luminescence were measured after 10 minutes incubation with each reagent respectively as per manufacturer’s instructions for the Dual-Glo Luciferase Assay System E2920 (Promega) using the Glomax luminometer (Promega).

#### IAV polymerase purification and *in vitro* activity assay

Influenza WSN (H1N1) recombinant polymerases were purified from HEK293T cells. Ten cm plates of adherent cells were transfected with 3 µg of pcDNA3-PB1, pcDNA3-PB2-TAP and pcDNA3-PA with PEI. Forty-eight hours post-transfection, cells were harvested in PBS and lysed on ice for 10 min in 500 µl lysis buffer (50 mM hepes pH 8.0, 200 mM NaCl, 25% glycerol, 0.5% Igepal CA-630, 1mM β-mercaptoethanol, 1x PMSF, 1x Protease Inhibitor cocktail tablet (Roche). Lysates were cleared by centrifugation at 17000 g for 5 min at 4°C, diluted in 2ml NaCl, and bound to pre-washed IgG sepharose beads for 2 h at 4°C. Beads were pre-washed 3x in binding buffer (10mM Hepes pH 8.0, 0.15 M NaCl, 0.1% Igepal CA-630, 10% glycerol, 1x PMSF). After binding, beads were washed 3x in binding buffer and 1x in cleavage buffer (10mM Hepes pH 8.0, 0.15M NaCl, 0.1% Igepal CA-630, 10% glycerol, 1x PMSF, 1mM DTT). Beads were cleaved with AcTEV protease overnight at 4°C, and cleared by centrifugation at 1000 g for 1 min as described previously(*33*).

#### Cell fractionation

Fractionation of transfected cells into cytoplasmic, mitochondrial, and nuclear components was carried out using the Abcam Cell Fractionation Kit (Abcam) following the manufacturer’s instructions, with volumes adjusted based on the number of cells. Samples of unfractionated whole cells in Buffer A were retained as input controls. Whole cells and sub-cellular fractions were dissolved in Trizol, for RNA extraction and analyzed as described above, or in 10%-SDS protein-loading buffer, for protein expression analysis by SDS-PAGE and western blot.

## Statistical testing

Statistical testing was carried out using GraphPad Prism 9 software. Error bars represent standard deviations, and either individual data or group mean values are plotted. One-way analysis of variance (ANOVA) with Dunnett’s test for multiple comparisons was used to compare multiple-group means to a normalized mean (e.g., IFN induction or RNA template replication). Two-way ANOVA with Sidak’s test for multiple comparisons was used to compare multiple pairs of group means (e.g., between two cell types, HEK293 WT to HEK293 MAVS^−/−^).

