## Supplemental Figure for "Transient RNA structures cause aberrant influenza virus replication and innate immune activation"

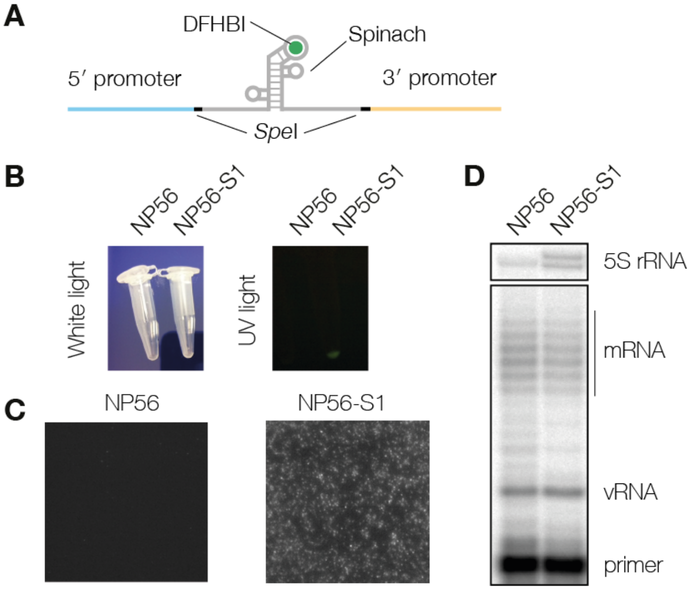


**Figure S1**. Effect of a single spinach aptamer on mvRNA replication. A) To rule out that a single RNA structure upstream of the RNA polymerase reduces activity, we engineered the NP56 mvRNA template and inserted the RNA aptamer Spinach, creating NP56-S1. B) A Spinach-containing mvRNAs generated in vitro demonstrated fluorescence after addition of DFHBI relative to the 56-nt control mvRNA, confirming that the aptamer folds properly in the context of the IAV promoter sequence. C) TIRF microscopy image of a 56-nt mvRNA or Spinach-containing mvRNA generated in vitro in the presence of DFHBI. D) After transfection of plasmids encoding the Spinach-containing mvRNA NP56-S1 into HEK293T cells and analysis of the RNA produced using primer extension, no difference in replication was observed relative to our NP56 control mvRNA.

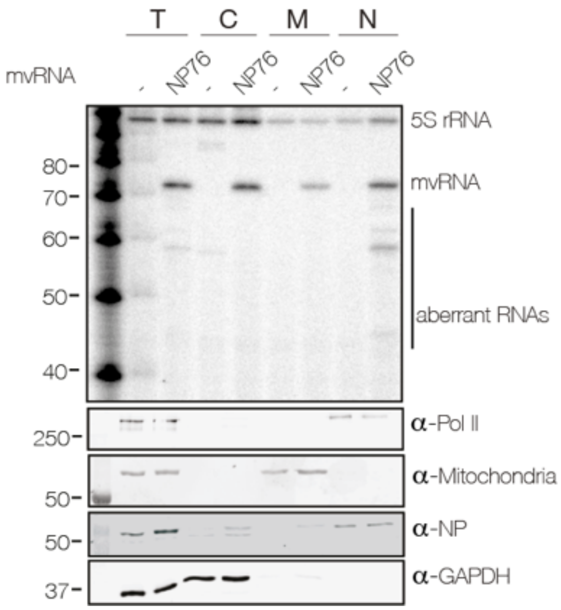


**Figure S2**. Fractionation of HEK 293T cells transfected with IAV RNA polymerase and a 76 nt-long segment 5-derived mvRNA. Top panel shows primer extension analysis of RNA extracted from fractions. Bottom four panels show western blot analysis. For each lane, comparable amounts were loaded.

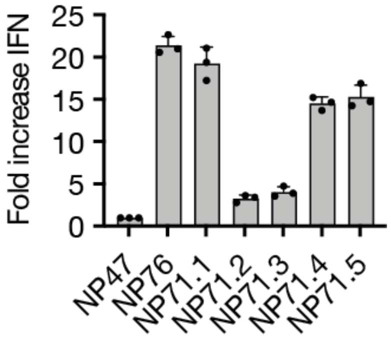


**Figure S3**. IFN-β promoter activity induced by the retransfection of total RNA isolated from HEK 293T cells transfected with plasmids expressing segment 5-derived mvRNAs and the viral RNA polymerase subunits. Luciferase signal was normalized to the NP47 mvRNA, which does not trigger innate immune responses.

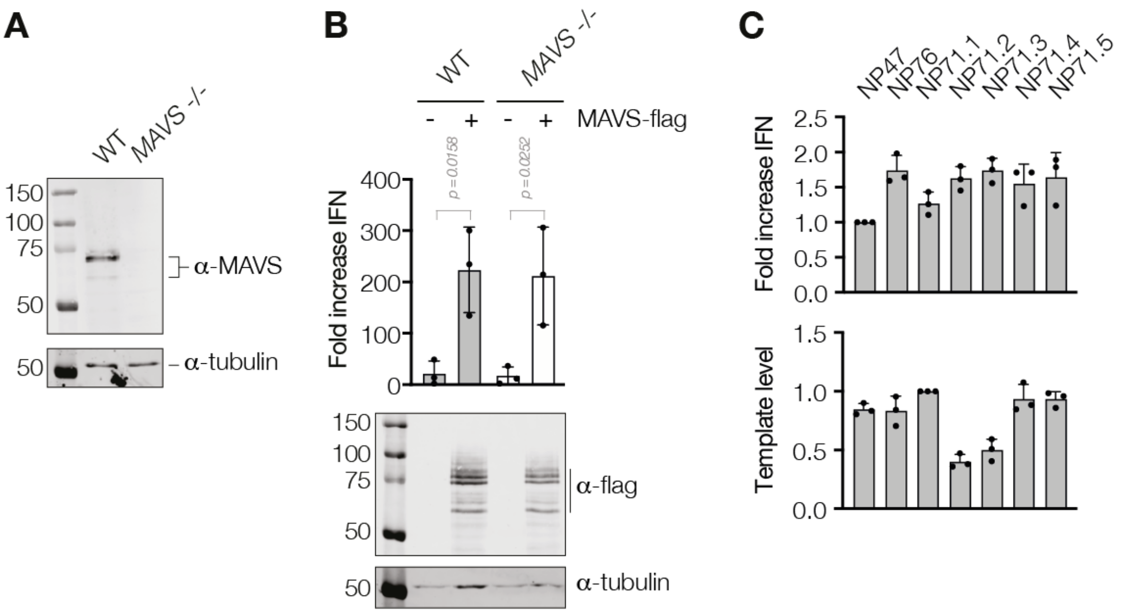


**Figure S4**. Characterization of *MAVS*$^{-/-}$ HEK 293 cells and mvRNA replication in *MAVS*$^{-/-}$ HEK 293 cells. A) Western blot analysis of MAVS expression in wildtype or *MAVS*$^{-/-}$ HEK 293 cells. B) IFN-β promoter activity in wildtype or *MAVS*$^{-/-}$ HEK 293 cells following transfection of a plasmid expressing MAVS-flag. Bottom panel shows western blot of MAVS-flag expression. C) IFN-β promoter activity of HEK 293 *MAVS*$^{-/-}$ cells expressing mvRNAs NP71.1-NP71.5. and primer extension analysis of mvRNA NP71.1-NP71.5 replication in HEK 293 *MAVS*$^{-/-}$ cells.

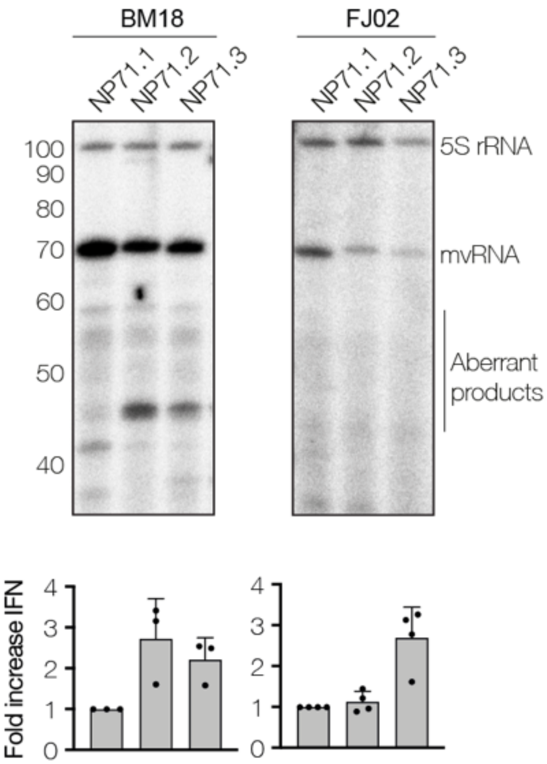


**Figure S5**. Replication of model mvRNAs in HEK 293T cells by the IAV A/Brevig Mission/1/18 (BM18) or A/duck/Fujian/01/02 (FJ02) RNA polymerases. The ability of these reactions to induce IFN-β promoter activity was analyzed using a luciferase reporter assay.

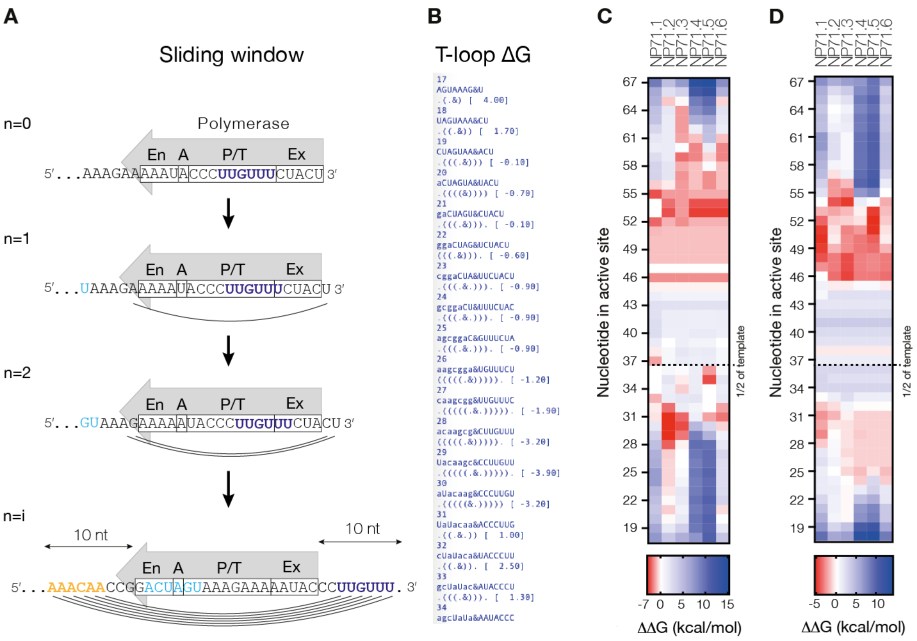


**Figure S6**. Sliding-window analysis of the free energy (ΔG) of transient secondary RNA structure formation near the influenza A virus RNA polymerase. A) To calculate the ΔG for a t-loop, 10 nt on either side of the polymerase were allowed to fold using the duplex-fold algorithm of the ViennaRNA package. To calculate the ΔG for alternative structures, cofold of the ViennaRNA package was used. The ΔΔG for a t-loop was calculated by subtracting the upstream and downstream ΔG values. B) Example of Python script output. C) ΔΔG values for positive sense 71-nt long mvRNA templates. D) ΔΔG values for negative sense 71-nt long mvRNA templates.

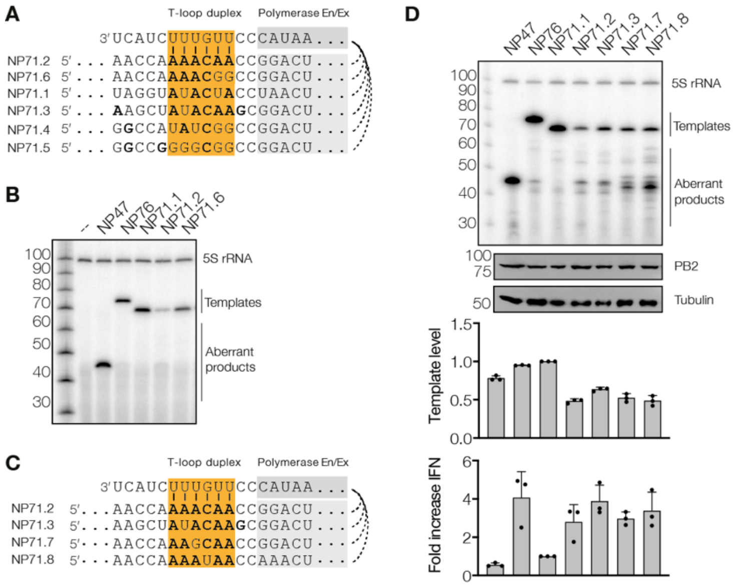


**Figure S7**. Replication of engineered mvRNAs and their ability to induce IFN-β promoter activity. A) Alignment of t-loops (shaded orange) in NP71 mvRNA templates. Location of RNA polymerase entry (En) and exit (Ex) channel is shaded gray. B) Primer extension analysis of RNA extracted from HEK 293T cells expressing NP71 mvRNA templates. C) Alignment of t-loops (shaded orange) of additional NP71 mvRNA templates. Location of RNA polymerase entry and exit channel is shaded gray. D) Primer extension analysis of RNA extracted from HEK 293T cells expressing NP71 mvRNA templates. Second and third panel show western blot analysis. Top graph shows quantification of mvRNA template level, while bottom graph shows IFN-β promoter activity analysis using a luciferase reporter.

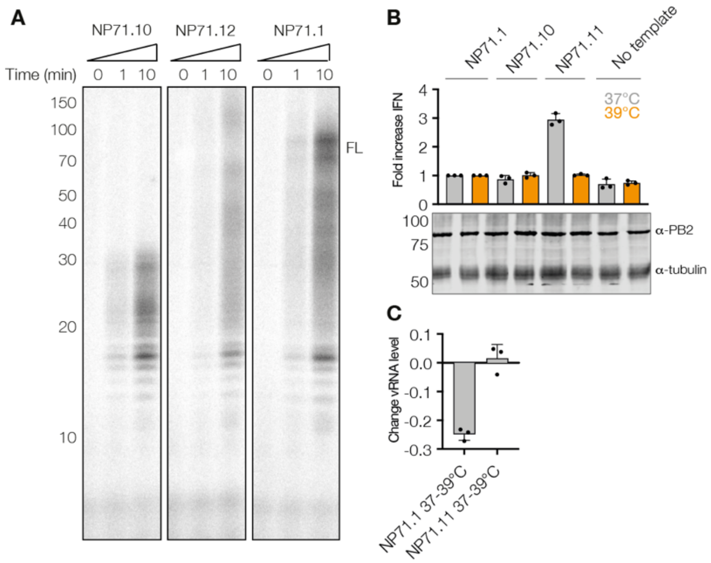


**Figure S8**. Effect of temperature on mvRNA template replication. A) Analysis of IAV RNA polymerase activity in vitro. B) Effect of temperature increase on viral replication and induction of IFN-β promoter activity. Graph shows fold increase IFN-β promoter activity. Bottom panel shows western blot analysis of viral protein expression. C) Quantification of vRNA level difference between incubations at 37 and 39 degrees Celsius for the NP71.1 and NP71.11 templates.

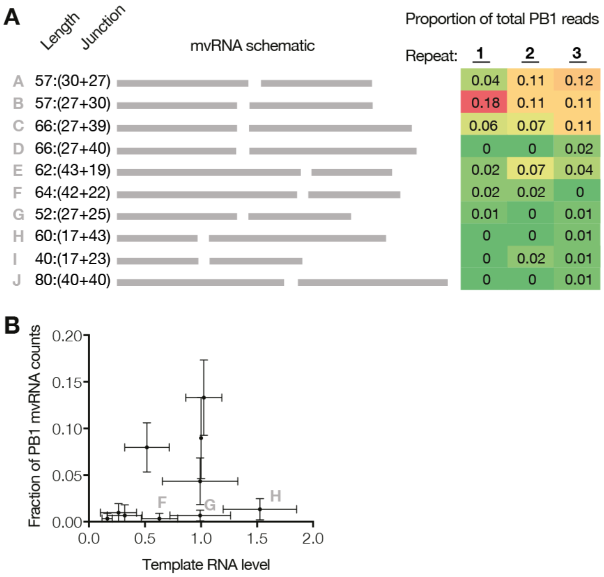


**Figure S9**. Selection of segment 2 mvRNAs. A) Segment 2 mvRNA sizes and abundance in three next generation sequencing experiments. B) Relation between mvRNA level in transfection assay and mvRNA read counts in next generation sequencing. The mvRNAs F, G and H are indicated.

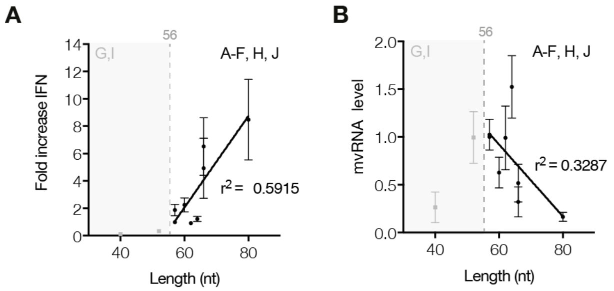


**Figure S10**. Correlation between the A) mvRNA template length and the IFN-β promoter activity induction, or B) the mvRNA template length and mvRNA replication.

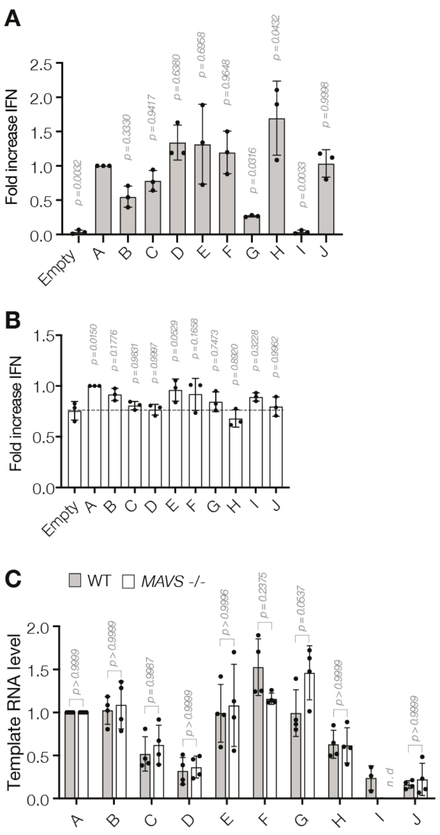

**Figure S11**. Segment 2 mvRNA replication in *MAVS*$^{-/-}$ HEK 293 cells. A) IFN-β promoter activity measured after retransfection of total RNA extracted from HEK 293T cells expressing segment 2 mvRNAs into HEK 293T cells. B) IFN-β promoter activity induced by expression of segment 2 mvRNAs in *MAVS*$^{-/-}$ HEK 293 cells. C) Primer extension analysis of segment 2 mvRNAs levels in wildtype and *MAVS*$^{-/-}$ HEK 293 cells.

**Table S1**. Sequences of mvRNA templates used.

| Name | Internal lab reference | Sequence 5′ and 3′ (vRNA) |
|---|---|---|
| NP71.1 | GC33 | AGUAGAAACAAGGGUAUUUUUCUUUACUAGUUAGGUAGUAUA CCUAGUAACUAGUCUACCCUGCUUUUGCU |
| NP71.2 | GC50.1 | AGUAGAAACAAGGGUAUUUUUCUUUACUAGUCCGGUUGUUUU GGUUGCCACUAGUCUACCCUGCUUUUGCU |
| NP71.3 | GC50.2 | AGUAGAAACAAGGGUAUUUUUCUUUACUAGUCCGCUUGUAUA GCUUGCCACUAGUCUACCCUGCUUUUGCU |
| NP71.4 | GC67 | AGUAGAAACAAGGGUAUUUUUCUUUACUAGUCCGGCCGAUAU GGCCGCCACUAGUCUACCCUGCUUUUGCU |
| NP71.5 | GC83 | AGUAGAAACAAGGGUAUUUUUCUUUACUAGUCCGGCCGCCCC GGCCGCCACUAGUCUACCCUGCUUUUGCU |
| NP71.6 | GC50.9 | AGUAGAAACAAGGGUAUUUUUCUUUACUAGUCCGGCCGUUUU GGUUGCCACUAGUCUACCCUGCUUUUGCU |
| NP71.7 | GC50.3 | AGUAGAAACAAGGGUAUUUUUCUUUACUAGUCCGGUUCUUUU GGUUGCCACUAGUCUACCCUGCUUUUGCU |
| NP71.8 | GC50.4 | AGUAGAAACAAGGGUAUUUUUCUUUACUAGUCCGGUUGCUUU GGUUGCCACUAGUCUACCCUGCUUUUGCU |
| NP47 | NP47 | AGUAGAAACAAGGGUAUUUUUCUUUACUAGUCUACCCUGCUU UUGCU |
| NP56 | NP56 | AGUAGAAACAAGGGUAUUUUUCUUUCUCGAGCGUACUAGUCU ACCCUGCUUUUGCU |
| NP76 | NP76 | AGUAGAAACAAGGGUAUUUUUCUUUACUAGUGAUUUCGAUGU CACUCUGUGAGUGAUUAUCUACCCUGCUUUUGCU |
| NP71.10 | GC50_13 | AGUAGAAACAAGGGUAUUUUUCUUUACUAGUGGCAGCAAAAG CAGGGUAACUAGUCUACCCUGCUUUUGCU |
| NP71.11 | GC50_15 | AGUAGAAACAAGGGUAUUUUUCUUUACUAGUGGCAGCAAAAG CACCCAUACUAGUCUACCCUGCUUUUGCU |
| NP71.12 | GC50_16 | AGUAGAAACAAGGGUAUUUUUCUUUACUAGUGGCUCUAAAAG CACCCAUACUAGUCUACCCUGCUUUUGCU |

**Table S2**. Cloned PB1 WSN mvRNAs.

| Name | Length (nt) | Sequence 5′ and 3′ (vRNA) |
|---|---|---|
| PB1 A | 57 | AGUAGAAACAAGGCAUUUUUUCAUGAAAUCCAUUCAAAUGGUUUGCCUGCUUUCGCU |
| PB1 B | 57 | AGUAGAAACAAGGCAUUUUUUCAUGAAGGACAUUCAAAUGGUUUGCCUGCUUUCGCU |
| PB1 C | 66 | AGUAGAAACAAGGCAUUUUUUCAUGAAGGACAAGCUAAACAUUCAAAUGGUUUGCCUGCUUUCGCU |
| PB1 D | 67 | AGUAGAAACAAGGCAUUUUUUCAUGAAGGACAAGCUAAAUCAUUCAAAUGGUUUGCCUGCUUUCGCU |
| PB1 E | 62 | AGUAGAAACAAGGCAUUUUAAGUCGGAUUGACAUCCAUUCAAAUGGUUUGCCUGCUUUCGCU |
| PB1 F | 64 | AGUAGAAACAAGGCAUUUUUUCAGUCGGAUUGACAUCCAUUCAAAUGGUUUGCCUGCUUUCGCU |
| PB1 G | 52 | AGUAGAAACAAGGCAUUUUUUCAUGCAUUCAAAUGGUUUGCCUGCUUUCGCU |
| PB1 H | 60 | AGUAGAAACAAGGCAUUUUUUCAUGAAGGACAAGCUAAAUUCAGUUUGCCUGCUUUCGCU |
| PB1 I | 40 | AGUAGAAACAAGGCAUUUUUUCAGUUUGCCUGCUUUCGCU |
| PB1 J | 80 | AGUAGAAACAAGGCAUUUUUUCAUGAAGGACAAGCUAAAUUCGGAUUGACAUCCAUUCAAAUGGUUUGCCUGCUUUCGCU |
